# Differential cortical and subcortical visual processing with eyes shut

**DOI:** 10.1101/2023.09.11.557197

**Authors:** Nicholas G. Cicero, Michaela Klimova, Laura D. Lewis, Sam Ling

## Abstract

Closing our eyes largely shuts down our ability to see. That said, our eyelids still pass some light, allowing our visual system to coarsely process information about visual scenes, such as changes in luminance. However, the specific impact of eye closure on processing within the early visual system remains largely unknown. To understand how visual processing is modulated when eyes are shut, we used functional magnetic resonance imaging (fMRI) to measure responses to a flickering visual stimulus at high (100%) and low (10%) temporal contrasts, while participants viewed the stimuli with their eyes open or closed. Interestingly, we discovered that eye closure produced a qualitatively distinct pattern of effects across the visual thalamus and visual cortex. We found that with eyes open, low temporal contrast stimuli produced smaller responses, across the lateral geniculate nucleus (LGN), primary (V1) and extrastriate visual cortex (V2). However, with eyes closed, we discovered that the LGN and V1 maintained similar BOLD responses as the eyes open condition, despite the suppressed visual input through the eyelid. In contrast, V2 and V3 had strongly attenuated BOLD response when eyes were closed, regardless of temporal contrast. Our findings reveal a qualitative distinct pattern of visual processing when the eyes are closed – one that is not simply an overall attenuation, but rather reflects distinct responses across visual thalamocortical networks, wherein the earliest stages of processing preserves information about stimuli but is then gated off downstream in visual cortex.

**Significance statement:** When we close our eyes, not all information is blocked out. Coarse luminance information is still accessible for processing by the visual system, even when our eyes are closed. Using functional magnetic resonance imaging (fMRI), we examined whether eyelid closure plays a unique role in visual processing. We discovered that while the thalamus and primary visual cortex (V1) show equivalent luminance-dependent responses both when the eyes are open and closed, extrastriate cortex exhibited a qualitatively distinct pattern of responses. Specifically, eye closure attenuated luminance responses in extrastriate cortices, but responses were preserved in LGN and V1. This pattern suggests that during brain states where the eyes are closed, visual information is still accessible to the very earliest stages of visual processing, but that downstream visual processing areas appear to become blind to this information.

## Introduction

Light exposure during sleep has substantial effects on the brain: it can alter circadian rhythms, sleep quality, and mood (Blume et al., 2019; Ohayon & Milesi, 2016). During sleep, our eyes are closed and the eyelids function as potent filters of visual information. However, our eyelids are only partial filters and do not completely attenuate all visual information (Ando & Kripke, 1996; Bierman et al., 2011). The eyelid has been characterized as a red-pass filter, with an estimated 6% red light spectral transmittance (Ando & Kripke, 1996). Indeed, subjective experience with high luminance stimuli, such as during a sunny day, corroborates the idea that changes in luminance are still detectable when our eyes are closed. With partial, rather than complete, filtering properties, it follows that the visual system processes external visual information with our eyes closed, as well.

How does the visual system process information when our eyes are closed? It is possible that the filtering properties of the eyelid simply quantitatively suppress responses across visual regions, due to the attenuation of input. Alternatively, eye closure could induce qualitatively distinct changes in visual response, selectively modulating responses in specific brain networks. While little is known about stimulus-evoked visual responses with eyes closed, resting-state fMRI studies have investigated spontaneous dynamics during eye closure in the absence of any visual stimulus presentation (Marx et al., 2003; Wei et al., 2018; Weng et al., 2020). These studies found differences in resting-state functional connectivity in attentional networks depending on whether eyes were open or closed, along with differences in activation in prefrontal cortex, parietal and frontal eye fields, and LGN. While eye closure appears to play a unique role in modulating brain responses, the impact that eye closure has on stimulus-evoked visual responses remains poorly understood.

In this study, we sought to shed light on the role that eye closure plays in modulating responses within the visual processing hierarchy. To do so, we measured fMRI BOLD responses within visual cortex and subcortex while participants viewed high and low intensity visual stimuli, with their eyes open or shut. We manipulated the intensity of visual input via temporal contrast modulation, in which the luminance of a uniform visual stimuli flickered rapidly between extreme whites and blacks (high temporal contrast), or between middling intensities (low temporal contrast). Indeed, previous work has shown visuocortical responses to be sensitive to changes in luminance (Vinke & Ling, 2020). By measuring BOLD responses to high and low luminance contrast stimuli, we examined whether there is a qualitatively unique pattern of luminance responses across the visuocortical hierarchy when one’s eyes are closed, compared to when they are open.

## Methods

### Participants

Data was acquired from a total of 8 healthy participants (5 females, 3 males). Participants were aged 18-35 years, reported normal or corrected-to-normal visual acuity, and were recruited from Boston University and the surrounding community. All participants provided written informed consent before study enrollment and completed a metal screening form indicating that they had no MRI contraindications. Participants were reimbursed for their study participation. All aspects of the study were approved by Boston University’s Institutional Review Board.

### Apparatus & stimuli

Stimuli were generated using custom software written in MATLAB (version 2019b) in conjunction with Psychtoolbox (Brainard, 1997). Participants viewed stimuli that was back-projected onto a screen set within the MRI scanner, using a ProPIXX DLP LED (VPixx Technologies) projector system (minimum luminance: 1.2 cd/m2; maximum luminance: 2507.9 cd/m^2^). Photometer measurements (model LS-100; Konica Minolta) carried out before the study were used to verify the linearity of the display (1 digital-to-analog conversion (DAC) step = 9.835 cd/m^2^). These measurements were used to calculate the stimulus luminance and were acquired from the inner-facing side of the back-projection screen while positioned within the MRI scanner bore. This was done to best account for the attenuation in luminance due to back-projection screen characteristics.

During each functional run, participants fixated on a median luminance crosshair at the center of the display while shown a full screen flickering display (17 degrees of visual angle) with no spatial contrast (Figure 1). The full field flicker was presented in a block design with three trial types (baseline, high, and low temporal luminance contrast), with each event lasting 16 seconds. In the *baseline* events, the full field display was a constant median luminance with no luminance modulation. During *high* events, the full field display flickered with an amplitude envelope of 100% around the middle luminance value. For *low* events the full field display flickered with an amplitude envelope of 10% around the median luminance value. All high and low events flickered at a frequency of 7 Hz.

**Figure 1.**
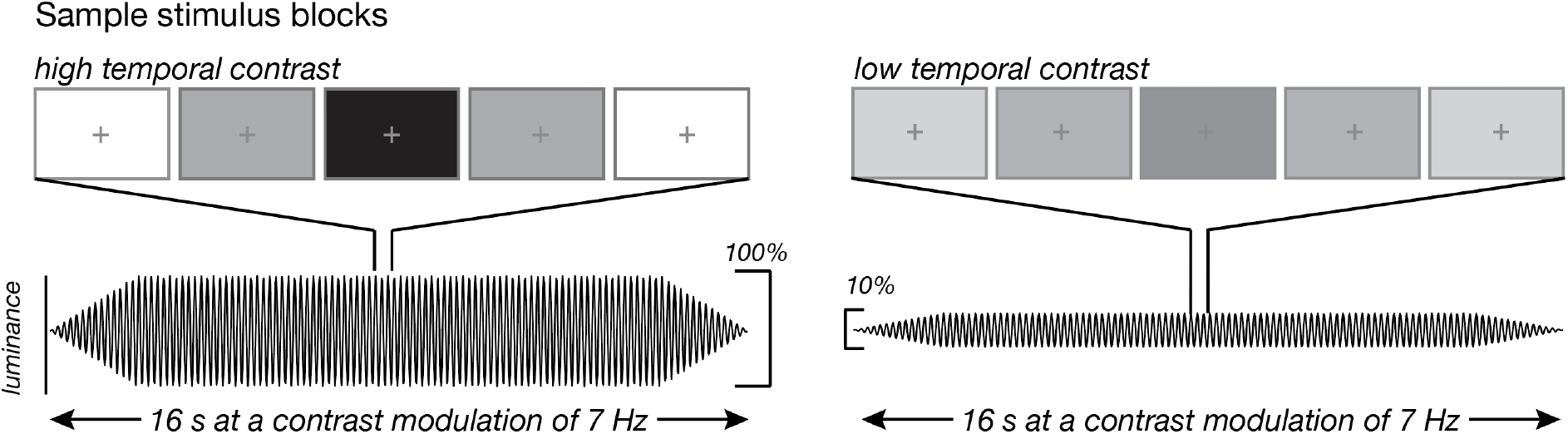
Experimental design with sample stimulus frames displaying the high temporal contrast and low temporal contrast displays. High temporal contrast flickered at 7 Hz with a luminance amplitude envelope of 100%, encompassing the maximum (255 a.u.) and minimum (0 a.u.) possible luminance values. The low temporal contrast events also flickered at 7 Hz with a luminance amplitude envelope of 10%, encompassing a range of luminance values between 140 a.u. and 115 a.u.

### Experimental design

Subjects participated in two scan sessions, each lasting approximately two hours. The first session was dedicated to collecting anatomical images and data for population receptive field (pRF) mapping using standard techniques and stimuli (Dumoulin & Wandell, 2008; Kay et al., 2013). The second session was dedicated to collecting proton-density (PD) weighted anatomical imaging and fMRI blood oxygenation level-dependent (BOLD) data across the eyes open and closed conditions, during the luminance task.

During the second experimental session, we collected three PD-weighted anatomical scans. PD-weighted anatomical imaging has previously been used to better localize the LGN (Fujita et al., 2001; Ling et al., 2015). Following the PD-weighted scans, participants completed three consecutive runs of a functional localizer. The visual stimulus for the functional localizer contained a full field flickering grating stimulus (diameter = 6.0°) with a centered circle (diameter = 0.8°). Within the centered circle, letters rapidly appeared one at a time with a new letter appearing every 200 ms. Participants were instructed to press a button whenever the letters ‘J’ and ‘K’ appeared within the centered circle. During the localizer blocks, the full field display alternated between a flickering grating stimulus and a full field non-flickering display at median luminance value. Participants completed 12 total blocks (6 flickering field, 6 non-flickering field) with an extra non-flickering block at the beginning of the run. At the end of each localizer run, participants were asked to report their wakefulness level.

Participants then completed the luminance flicker task. The task began and ended with a baseline event. High and low temporal contrast conditions were pseudo-randomly ordered, with all high and low events interleaved with a baseline event. Each run contained 12 events (6 high, 6 low) interspersed with 12 baseline events, lasting a total of 384 seconds. On each run participants were instructed to press a button after each full breath cycle (1 inhale, 1 exhale). This button task was chosen to ensure that participants did not fall asleep and engaged with the task, while not requiring eyes to be open. For each run, participants were instructed to either keep their eyes open and fixate on the crosshair or to keep their eyes closed throughout the run. Each scan session began with an eyes-closed run, and consecutive runs alternated between open and closed conditions. We always began with the eyes closed condition to ensure we acquired a sufficient number of runs in this condition, where BOLD modulations may be lower compared to eyes-open runs. To ensure participants kept their eyes closed or open, real time eye monitoring was carried out using an EyeLink1000, for the duration of each run. On average, we collected 5 runs with eyes closed and 4 runs with eyes open, for each subject.

### MRI data acquisition

All neuroimaging data were acquired using a research-dedicated Siemens Prisma 3T scanner using a Siemens 64-channel head coil. A whole brain anatomical scan was acquired using a T1-weighted multi-echo MPRAGE (1 mm isotropic voxels; field of view (FOV) = 192 × 192 × 134 mm, flip angle (FA) = 7.00°, repetition time (TR) = 2200 ms, echo time (TE) = 1.57 ms). Proton density (PD)-weighted anatomical scans were acquired to localize LGN (0.9mm × 0.9mm × 1.7mm; TR = 2950.0 ms; TE = 15.6 ms; FA = 180°). Functional scans were acquired using T2*-weighted in-plane simultaneous imaging (2 mm isotropic voxels; FOV = 104 × 104 × 70 mm, FA = 64.00°, TR = 1000 ms, TE = 30 ms, SMS factor = 5, GRAPPA acceleration = 2).

### Anatomical data analysis

T1-weighted anatomical data were analyzed using the standard “recon-all” pipeline provided by the FreeSurfer neuroimaging analysis package (Fischl, 2012), generating cortical surface models, whole brain segmentations, and cortical parcellations. All PD-weighted scans were aligned to each subject’s anatomical space and averaged together (using AFNI’s 3dcalc).

### Functional data analysis

Functional BOLD time-series data were first corrected for echo-planar imaging (EPI) distortions using a reverse phase-encode method implemented in FSL (Andersson et al., 2003) and were then preprocessed with FS-FAST using standard motion-correction procedures, slice timing correction, and boundary-based registration between functional and anatomical spaces (Greve & Fischl, 2009). To optimize spatial precision of experimental data, no volumetric spatial smoothing was performed (full-width half-maximum 0 mm). To achieve precise alignment of experimental data within the session, cross-run within-modality robust rigid registration was performed, using the middle time point of each run (Reuter et al., 2010). BOLD time-series data were demeaned and converted to units of percent signal change. Data collected during the separate pRF mapping scans were analyzed using the analyzePRF toolbox (Kay et al., 2013). Results from the pRF model were used to manually draw labels for our regions of interest within visual cortex.

### Statistical analysis

The results from the pRF modeling were used to identify region-of-interest (ROI) labels for each cortical region before analysis. ROI labels included voxels located inside the cortical ribbon for V1/V2/V3, which were identified using a visual area network label generated using an intrinsic functional connectivity atlas (Yeo et al, 2012). Results from the pRF modeling were additionally used to select voxels with visual field eccentricity preferences less than 17 degrees visual angle away from fixation as this was the measured extent of the screen within the MRI scanner. Cortical voxels with a poor pRF model fit (r^2^ < 0.10) were removed from further analyses. Initial LGN labels were acquired from thalamic segmentation and parcellation in Free-Surfer for each participant. These initial labels were overlaid with the GLM results from the functional localizer and the PD-weighted scans, and only intersecting voxels from the top 40% of t-values from the functional localizer were chosen for the final LGN labels and further analyses.

An event-triggered average was computed for each flickering condition (low and high) per eyelid condition and ROI. The BOLD time-series for each ROI per run was separated by the low and high trials, and all trials of a given type were averaged together. Average BOLD magnitude in response to the stimulus presentation was computed by averaging 4-16 s post-stimulus onset for each trial. Two-way between-subjects ANOVA were performed to test for any main effects of temporal contrast and eye closure and any interaction of the two on average BOLD magnitude during stimulus presentation. Additional event-triggered average analysis was done with eccentricity, in which the time-series for V1/V2/V3 voxels were first separated into eccentricity bins defined by degree visual angle relative to fixation. Foveal-tuned voxels were between 0.01° – 1.5°, parafoveal-tuned voxels were between 1.5° – 4.0°, and peripheral-tuned voxels were between 4.0° – 17.0°. An additional ANOVA was performed to test for any main effect of eccentricity on BOLD response during stimulus presentation. Multiple comparison correction was done using Bonferroni correction of α/n at a familywise α of 0.05 where n is the number of tests performed.

## Results

We first examined how temporal contrast modulated thalamic and visuocortical responses, and if eye closure impacted these responses. With eyes open, LGN, V1, and V2 showed larger responses to high temporal contrast stimuli, compared to low temporal contrast stimuli (Figure 2A). Indeed, during eyes open with high temporal contrast stimuli, all ROIs had significantly elevated BOLD responses [LGN: *t*(7) = 2.84, *P* = 0.0125; V1: *t*(7) = 5.45, *P* < 0.0001; V2: *t*(7) = 3.92, *P* = 0.0028; V3: *t*(7) = 2.44, *P* = 0.022), though the significant response in V3 did not survive multiple comparisons correction. When the participants closed their eyes, however, LGN and V1 maintained their stronger responses to higher contrast stimuli [LGN: *F*(1,31) = 3.86, *P* = 0.059; V1: *F*(1,31) = 5.70, *P* = 0.023], which did not differ from their eyes closed conditions [LGN: *F*(1,31) = 0.847, *P* = 0.364; V1: *F*(1,31) = 3.06, *P* = 0.091]. In other words, while responses in LGN and V1 were significantly modulated by temporal contrast, they were completely unaffected by eye closure, despite the profound suppression of visual input from the eyelid.

**Figure 2.**
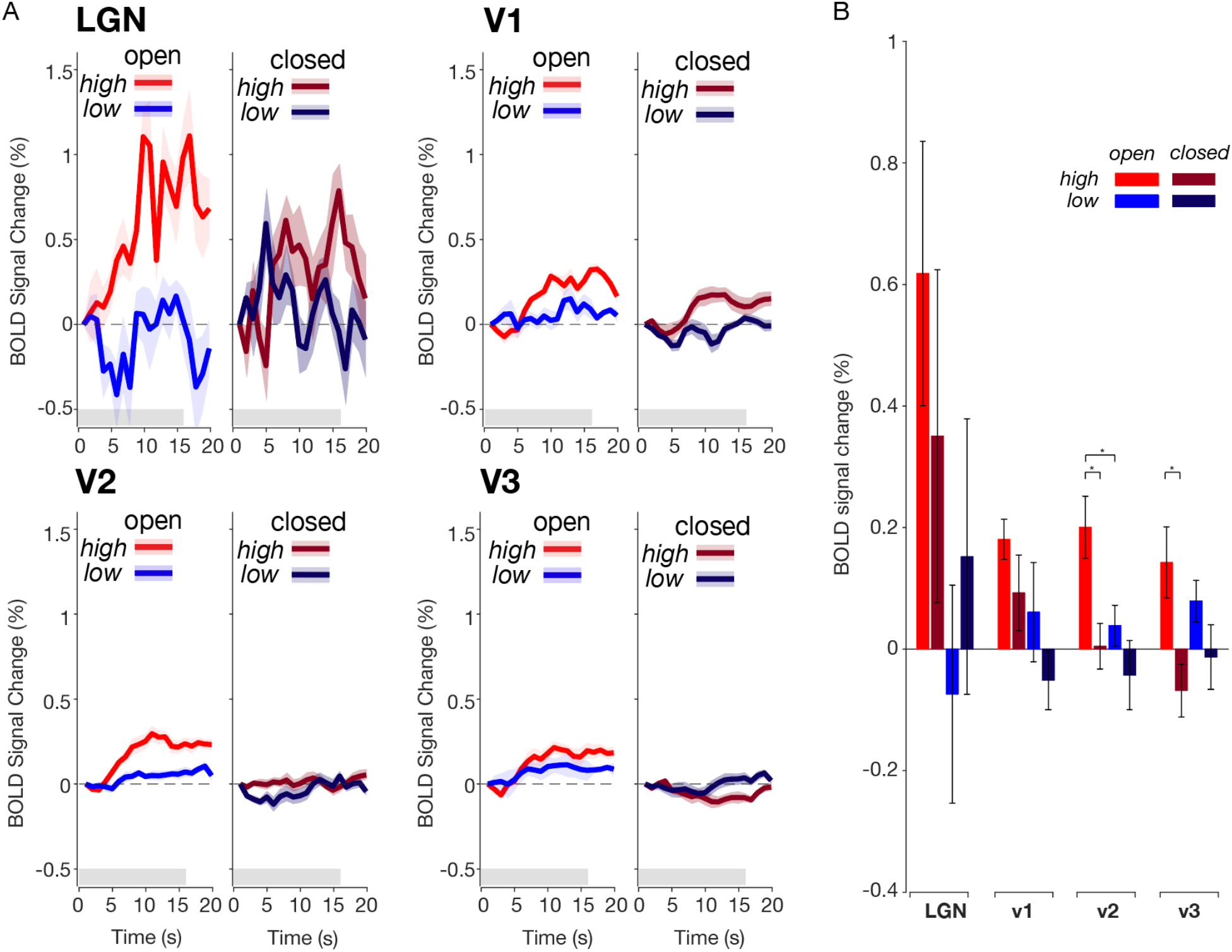
Eye closure has minimal effect on visual responses in LGN and V1, while suppressing responses in V2 and V3. (A) Event-triggered average for luminance task across ROI and eye condition. Across LGN, V1, and V2, during eyes open runs high temporal contrast stimuli elicits a greater BOLD response than with low temporal contrast stimuli. Though there is no effect of temporal contrast in V3, BOLD response increases regardless of the stimulus temporal contrast. During eye closure, BOLD responses in LGN and V1 during the high temporal contrast stimuli elicits a similar BOLD response as during eyes open runs. With eye closure, V2 and V3 have strongly attenuated BOLD regardless of temporal contrast. Red plots indicate high temporal contrast trials and blue indicates the low temporal contrast trials. The grey bar indicates 16 second period of stimulus presentation. Error shading is 1 SEM. N=8 subjects. (B) Average BOLD activation during stimulus presentation across conditions. Pairwise comparisons show a significant decrease in V2 and V3 BOLD magnitude with eye closure for high temporal contrast stimuli. In LGN, BOLD magnitude with high temporal contrast stimuli with eyes open was marginally greater than low contrast (*t*(14)=2.45; *P* = 0.0139) at a Bonferroni corrected p-value cutoff of 0.0125. In V1 with eyes closed, BOLD magnitude during high temporal contrast stimuli was marginally greater than during low temporal contrast stimuli (*t*(14)=1.82; *P* = 0.044). In V2, BOLD magnitude with high temporal contrast stimuli with eyes open was greater than low contrast (*t*(14)=2.65; *P* = 0.009) and high temporal contrast stimuli with eyes closed was suppressed compared to eyes open. In V3, BOLD magnitude during high temporal contrast stimuli with eyes closed was also suppressed compared to eyes open. Y-axis is BOLD signal averaged across 4-16s post-stimulus onset. Error bars are 1 SEM. All p-values from pairwise comparison only survive multiple comparison correction at a p-value less than 0.0125, using Bonferroni correction (0.05/n where n=4 per ROI). * *P* < 0.0125

Interestingly, while eye closure did not appear to have a major effect on the earliest stages of visual processing (LGN and V1), we observed a qualitatively distinct pattern within extrastriate cortices V2 and V3. When the eyes were closed, there was a drastic attenuation of stimulus evoked responses, regardless of temporal contrast [Main effects of eye closure: V2: *F*(1,31) = 9.66, *P* = 0.0043; V3: *F*(1,31) = 10.42, *P* = 0.003; Main effect of temporal contrast V2: *F*(1,31) = 5.54, *P* = 0.025; V3: *F*(1,31) = 0.01, *P* = 0.971]. Pairwise comparisons revealed a significant decrease in BOLD response to high temporal contrast stimuli with eye closure in V2 (*t*(14)=-3.09; *P* = 0.004) and V3 (*t*(14)=-2.89; *P* = 0.005). Overall, these results indicate that visual processing appears to be qualitatively different with eyes closed compared to when eyes are open. The BOLD response in LGN and V1 was modulated by temporal contrast but was unaffected by eye closure, whereas eye closure strongly reduced responses in extrastriate cortices V2 and V3.

Along with the heterogeneity in patterns observed across striate and extrastriate regions, it is possible that there exists heterogeneity *within* each region. It has been reported that there is an eccentricity bias of the BOLD response in V1 and V2, when participants viewed center-surround stimuli with no local contrast (Cornelissen et al., 2006). To test for an eccentricity bias and if eye closure impacts this bias, we separated voxels in V1-V3 by their eccentricity preference, based on pRF estimates (LGN was excluded from this analysis due to being underpowered for pRF analyses). We defined foveally-preferring voxels as those preferring between 0.01° – 1.5° from fixation, parafoveal-preferring voxels were those between 1.5° – 4.0°, and peripheral-preferring voxels were between 4.0° – 17.0°. As low temporal contrast trials elicited no significant activation across visuocortical regions, we did not test for an effect of eccentricity during low temporal contrast trials. We found that the effect of eccentricity was not significant in V1 [*F*(2,47) = 1.23, *P* = 0.303] (Figure 3), nor in V2 [*F*(2,47) = 0.90, *P* = 0.413] nor V3 [*F*(2,47) = 0.84, *P* = 0.440]. No ROIs had any significant interaction between eye closure and eccentricity [V1: *F*(2,47) = 1.23, *P* = 0.303; V2: *F*(2,47) = 0.47, *P* = 0.631; V3: *F*(2,47) = 0.48, *P* = 0.620]. This suggests that across striate and extrastriate cortices there is no eccentricity bias in BOLD responses nor any difference with eye closure. Thus, the impact of high temporal contrast stimuli and eye closure on BOLD appear uniform within each visuocortical area.

**Figure 3.**
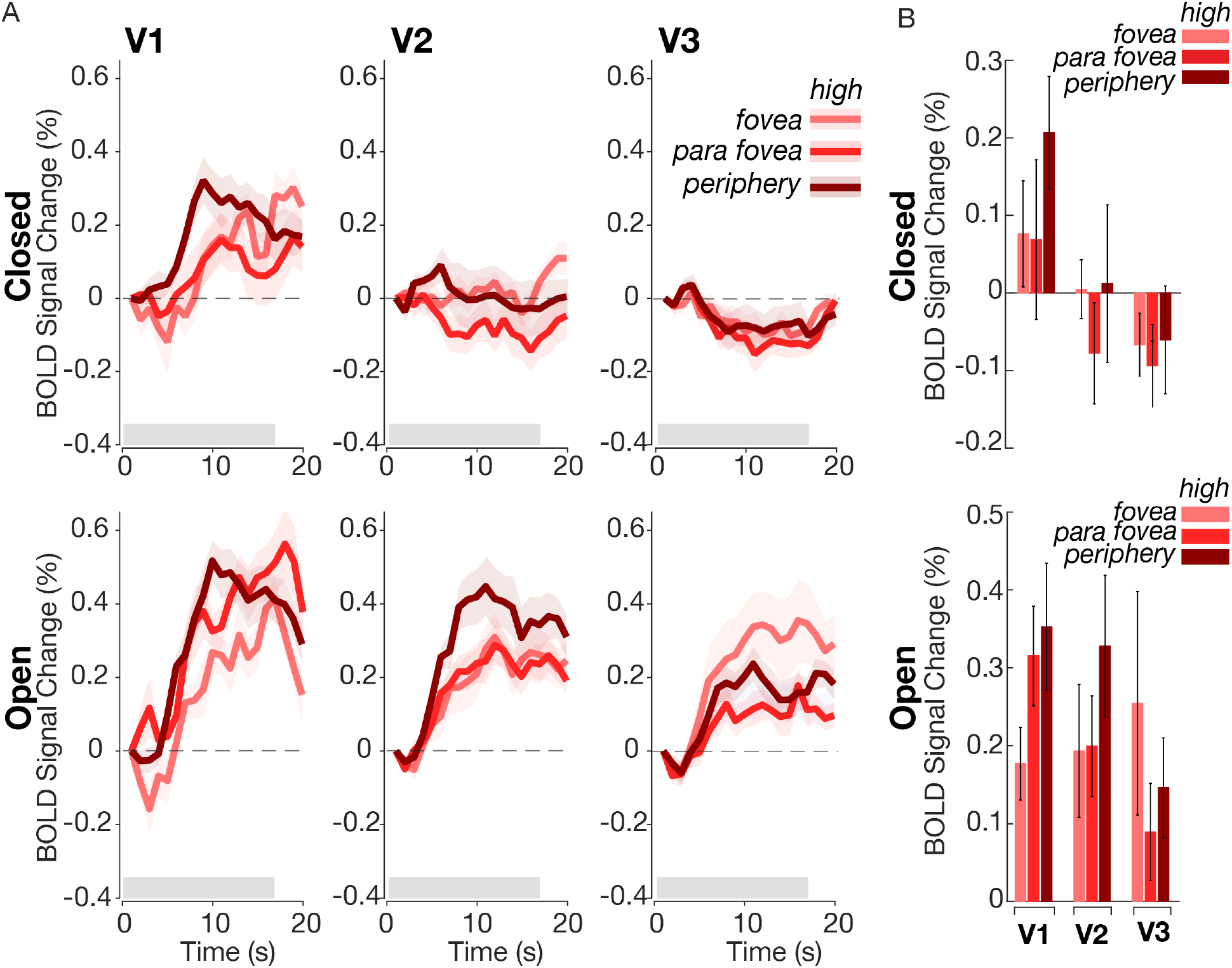
The effects of eye closure do not depend on eccentricity tuning. (A) Event-triggered average for BOLD response to luminance task across cortical ROI and eye condition separated by voxels tuned to different portions of the visual field. With eyes open and eyes closed, the BOLD responses to high contrast stimuli are uniform across eccentricities for all cortical ROIs. Foveal voxels were tuned to between 0.01 dva – 1.5 dva. Parafoveal voxels were tuned to between 1.5 dva – 4.0 dva. Peripheral voxels were tuned to between 4.0 dva – 17 dva. (B) Average BOLD activation during stimulus presentation across conditions (top = eyes closed; bottom = eyes open), separated by eccentricity preference. There are no significant pairwise comparisons when comparing eccentricity responses within each ROI. Y-axis is BOLD signal averaged across 4-16s post-stimulus onset. Error bars and error shading is 1 SEM.

## Discussion

With subjective experience it is clear that we can still perceive visual stimuli with closed eyes, but how distinct stages of the visual system supported this filtered visual experience was unknown. In this study, we found that eye closure produces a qualitatively distinct pattern of modulatory responses within the early visual system: closing one’s eyes selectively attenuated luminance processing in extrastriate cortex, but not in LGN nor striate cortex.

In line with previous literature showing that early visual responses can still occur when the eyes are closed (Marx et al., 2003; Sharon & Nir, 2018), we demonstrated that with closed eyes, luminance-dependent responses remain present in the LGN and V1. However, we found substantial heterogeneity in activation across regions when eyes were closed. One hypothesis as to why we observed strongly attenuated BOLD with closed eyes in extrastriate cortex, but not the LGN nor striate cortex, is that top-down modulation of visuocortical responses is often stronger in extrastriate compared to striate cortex (Haenny & Schiller, 1988; Moran & Desimone, 1985; Shulman et al., 1997). It has been demonstrated that higher-order sensory regions, such as the frontal eye field (FEF), may account for the selective top-down modulation of extrastriate cortical responses (Veniero et al., 2021). Resting-state fMRI studies that examined altered functional connectivity between eyes open and closed states found increased activation of the FEF during eyes closed relative to eyes open scans (Weng et al., 2020), lending further support to top-down modulation of extrastriate cortex during eyes closed states. Interestingly, one study which microstimulated the FEF of monkeys and measured visuocortical responses with fMRI found that FEF stimulation modulated extrastriate areas only in the presence of a visual stimulus, indicating that top-down modulation of the extrastriate cortices is dependent on bottom-up influence (Ekstrom et al., 2008). Since our paradigm includes a visual stimulus, it is possible that eye closure in the presence of visual stimuli attenuates extrastriate cortical responses through both top-down and bottom-up mechanisms. The eyelid abolishes almost all structure and form-like information, which is necessary to elicit responses in extrastriate cortices that prefer higher-level feature selectivity, such as spatial contrast, shapes, and contours. However, eyelid closure still passes through luminance information, which is known to activate striate cortex (Vinke & Ling, 2020). This preservation of luminance information, but attenuation of higher-level information, may explain the preservation of early visual pathway activation with weakened extrastriate activation.

Visuocortical responses have been shown to depend on luminance modulation, with responses increasing monotonically with luminance (Vinke & Ling, 2020). In addition to luminance modulation, luminance response functions are strongly contrast dependent, with lower spatial contrast drastically decreasing visuocortical responses to luminance (Vinke & Ling, 2020). Since the eyelid filters out much visual information, it is likely that spatial contrast no longer impacts visual responses and that luminance information dominates what might pass through the eyelid. Additionally, the lower luminance retinal input with eye closure cannot fully explain our results since LGN and V1 showed no significant change in BOLD activation between open and closed eye conditions. Since the eyelid is characterized as a red-pass filter (Ando & Kripke, 1996), it is possible that early visual pathways preferentially process this red visual content that extrastriate cortex is blind to; however, to our knowledge no evidence of this exists. Although further work will be needed to better unpack luminance responses in the early visual system, our results suggest that luminance-based responses within early visual areas may not always necessitate the existence of spatial contrast in order to reveal themselves, as previously suggested.

One interpretation of our results is that that modulation of visual processing during eye closure may be dependent on brain state, not just the physical barrier of the eyelid. Eye closure likely induces a change in overall brain state that alters both the processing of visual information and large-scale functional network processing. Eye closure decreases activity in attentional systems in the occipital and parietal lobes and increases functional coupling between sensory thalamus and somatosensory regions (Marx et al., 2003; Wei et al., 2018; Weng et al., 2020). These differences in spontaneous brain activity across sensory and attentional systems point to altered brain states with eye closure. Exteroceptive and interoceptive mental state hypotheses have been formulated where an exteroceptive mental state is characterized by increased attention and sensory processing of the external environment with eyes open (Marx et al., 2003). On the other hand, an interoceptive mental state is characterized by internally-directed cognition and reduced sensory processing with eyes closed. Many brain states require prolonged periods of eye closure, such as sleep and meditation, that involve reduced sensory awareness of external stimuli and enhanced internally-directed attention. Thus, eye closure may modulate visual processing through attentional or brain-state-dependent mechanisms.

## Author contributions

Conceptualization, N.G.C, M.K., L.D.L., and S.L.; Methodology, N.G.C., M.K., L.D.L., and S.L.; Investigation, N.G.C and M.K.; Writing – Original Draft, N.G.C, M.K., L.D.L., and S.L.; Writing – Review & Editing, N.G.C, M.K., L.D.L., and S.L.; Funding Acquisition, L.D.L. and S.L.; Supervision, L.D.L. and S.L.

Further information and requests for resources should be directed to and will be fulfilled by the lead contact, Sam Ling.

## Acknowledgements

This work was supported by National Institutes of Health R01 EY028163 to S.L., the Sloan Fellowship, McKnight Scholar Award, Pew Biomedical Scholar Award and NIH U19- NS128613 to L.D.L. This research was carried out at the Boston University Cognitive Neuroimaging Center. This work involved the use of instrumentation supported by the NSF Major Research Instrumentation grant BCS- 1625552. We acknowledge the University of Minnesota Center for Magnetic Resonance Research for use of the multiband-EPI pulse sequences. Data was analyzed on a high- performance computing cluster supported by the ONR grant N00014-17-1-2304. We thank Shruthi Chakrapani and Stephanie McMains for assistance with data collection, and members of the S.L. laboratory and L.D.L. laboratory for helpful feedback on the manuscript.

## References

Andersson, J. L. R., Skare, S., & Ashburner, J. (2003). How to correct susceptibility distortions in spin-echo echo-planar images: Application to diffusion tensor imaging. NeuroImage, 20(2), 870–888. 10.1016/S1053-8119(03)00336-7

Ando, K., & Kripke, D. F. (1996). Light Attenuation by the Human Eyelid.

Bierman, A., Figueiro, M. G., & Rea, M. S. (2011). Measuring and predicting eyelid spectral transmittance. Journal of Biomedical Optics, 16(6), 067011. 10.1117/1.3593151

Blume, C., Garbazza, C., & Spitschan, M. (2019). Effects of light on human circadian rhythms, sleep and mood. In Somnologie (Vol. 23, Issue 3, pp. 147–156). Dr. Dietrich Steinkopff Verlag GmbH and Co. KG. 10.1007/s11818-019-00215-x

Brainard, D. H. (1997). The Psychophysics Toolbox. Spatial Vision, 10(4), 433–436. 10.1163/156856897X00357

Cornelissen, F. W., Wade, A. R., Vladusich, T., Dougherty, R. F., & Wandell, B. A. (2006). No functional magnetic resonance imaging evidence for brightness and color filling-in in early human visual cortex. Journal of Neuroscience, 26, 3634–3641. 10.1523/JNEUROSCI.4382-05.2006

Dumoulin, S. O. & Wandell, B. A. (2008). Population receptive field estimates in human visual cortex. NeuroImage, 39, 647–660. 10.1016/j.neuroimage.2007.09.034

Ekstrom, L. B., Roelfsema, P. R., Arsenault, J. T., Bonmassar, G., & Vanduffel, W. (2008). Bottom-up dependent gating of frontal signals in early visual cortex. Science, 321, 414-417. doi: 10.1126/science.1153276

Fischl, B. (2012). FreeSurfer. Neuroimage 62:774–781.

Fujita, N., Tanaka, H., Takanashi, M., Hirabuki, N., Abe, K., Yoshimura, H., & Nakamura, H. (2001). Lateral geniculate nucleus: Anatomic and functional identification by use of MR imaging. American Journal of Neuroradiology, 22, 1719–1726. PMCID: PMC7974446

Greve, D. N., & Fischl, B. (2009) Accurate and robust brain image alignment using boundary-based registration. Neuroimage, 48, 63–72.

Haenny, P. E., & Schiller, P. H. (1988). State dependent activity in monkey visual cortex. Experimental Brain Research, 69(2), 225–244. 10.1007/BF00247569

Kay, K. N., Winawer, J., Mezer, A., & Wandell, B. A. (2013). Compressive spatial summation in human visual cortex. J Neurophysiol, 110, 481–494. 10.1152/jn.00105.2013.-Neurons

Ling, S., Pratte, M. S., & Tong, F. (2015). Attention alters orientation processing in the human lateral geniculate nucleus. Nature Neuroscience, 18(4), 496–498. 10.1038/nn.3967

Marx, E., Deutschländer, A., Stephan, T., Dieterich, M., Wiesmann, M., & Brandt, T. (2003). Eyes open and eyes closed as rest conditions: Impact on brain activation patterns. NeuroImage, 21(4), 1818–1824. 10.1016/j.neuroimage.2003.12.026

Moran, J., & Desimone, R. (1985). Selective Attention Gates Visual Processing in the Extrastriate Cortex. Science, 229(4715), 782–784. 10.1126/science.4023713

Ohayon, M. M., & Milesi, C. (2016). Artificial outdoor nighttime lights associate with altered sleep behavior in the American general population. Sleep, 39(6), 1311–1320. 10.5665/sleep.5860

Sharon, O. & Nir, Y. (2018). Attenuated fast steady-state visual evoked potentials during human sleep. Cerebral Cortex, 28, 1297–1311. doi:10.1093/cercor.bhx043

Shulman, G., Corbetta, M., Buckner, R. L., Raichle, M. E., Fiez, J. A., Miezin, F. M., & Petersen, S. E. (1997). Top-down modulation of early sensory cortex. Cerebral Cortex, 7(3), 193–206. 10.1093/cercor/7.3.193

Veniero, D., Gross, J., Morand, S., Duecker, F., Sack, A. T., & Thut, G. (2021). Top-down control of visual cortex by the frontal eye fields through oscillatory realignment. Nature Communications, 12(1). 10.1038/s41467-021-21979-7

Vinke, L. N., & Ling, S. (2020). Luminance potentiates human visuocortical responses. 2020. First Published De-Cember, 123, 473–483. 10.1152/jn.00589.2019.-Our

Wei, J., Chen, T., Li, C., Liu, G., Qiu, J., & Wei, D. (2018). Eyes-open and eyes-closed resting states with opposite brain activity in sensorimotor and occipital regions: Multidimensional evidences from machine learning perspective. Frontiers in Human Neuroscience, 12. 10.3389/fnhum.2018.00422

Weng, Y., Liu, X., Hu, H., Huang, H., Zheng, S., Chen, Q., Song, J., Cao, B., Wang, J., Wang, S., & Huang, R. (2020). Open eyes and closed eyes elicit different temporal properties of brain functional networks. NeuroImage, 222. 10.1016/j.neuroimage.2020.117230

Yeo, B. T., Krienen, F. M., Sepulcre, J., Sabuncu, M. R., Lashkari D., Hollinshead, M., Roffman, J. L., Smoller, J. W., Zöllei, L., Polimeni, J. R., Fischl, B., Liu, H., & Buckner, R. L. (2011) The organization of the human cerebral cortex estimated by intrinsic functional connectivity. J Neurophysiol, 106, 1125–1165.

